# Postural Representations of the Hand in Primate Sensorimotor Cortex

**DOI:** 10.1101/566539

**Authors:** James M. Goodman, Gregg A. Tabot, Alex S. Lee, Aneesha K. Suresh, Alexander T. Rajan, Nicholas G. Hatsopoulos, Sliman J. Bensmaia

## Abstract

Dexterous hand control requires not only a sophisticated motor system but also a sensory system to provide tactile and proprioceptive feedback. To date, the study of the neural basis of proprioception in cortex has focused primarily on reaching movements, at the expense of hand-specific behaviors such as grasp. To fill this gap, we record both the time-varying hand kinematics and the neural activity evoked in somatosensory and motor cortices as monkeys grasp a variety of different objects. We find that neurons in somatosensory cortex, as well as in motor cortex, preferentially track postures of multi-joint combinations spanning the entire hand. This contrasts with neural responses during reaching movements, which preferentially track movement kinematics of the arm rather than its postural configuration. These results suggest different representations of arm and hand movements likely adapted to suit the different functional roles of these two effectors.

## Introduction

Proprioception – the ability to track the postures and movements of our limbs – is critical to our ability to move fluidly and interact effectively with our environment, as evidenced by the devastating impairments experienced by individuals who have lost this sense (Brochier et al., 1999a; Cole, 2009; Cole and Paillard, 1998; Cole et al., 2002; Ghez et al., 1995; Kruger and Porter, 1958; Sainburg et al., 1995). Despite its importance, little is known about the cortical basis of proprioception, particularly that of the hand: Few proprioceptive studies have focused on representations of the digits (Costanzo and Gardner, 1981; Gardner and Costanzo, 1981), with the majority of work focusing on representations of the proximal limb (upper arm and forearm) (Fromm and Evarts, 1982; Fromm et al., 1984; London and Miller, 2013; Prud’homme and Kalaska, 1994; Weber et al., 2011). Fundamental differences exist between the hand and arm in terms of their respective functions and biomechanical properties: The arm is more massive and functions to transport the hand in three-dimensional space, whereas the hand is light and functions to conform to objects to enable grasp and manipulation of them. As such, the neural mechanisms that control and track these two effectors may be fundamentally different.

In the present study, we investigated the neuronal representations of hand postures and movements in somatosensory cortex during the most common manual activity of daily living, namely grasping. Specifically, we had monkeys grasp objects of varying shapes, sizes, and orientations – designed to elicit a wide variety of grasping movements – as we tracked the kinematics of the hand and measured neural activity in somatosensory cortex (SC) as well as primary motor cortex (M1) using chronically implanted electrode arrays (Figure 1) (Figure S1). In SC, we focused on cortical fields that are known to contain proprioceptive neurons, namely Brodmann’s areas 3a and 2, and verified electrode locations within these cortical fields with histology (Figure 1E). Our goal was to determine which aspects of hand movements and postures drive the responses of somatosensory neurons and contrast these to their motor counterparts.

**Figure 1.**
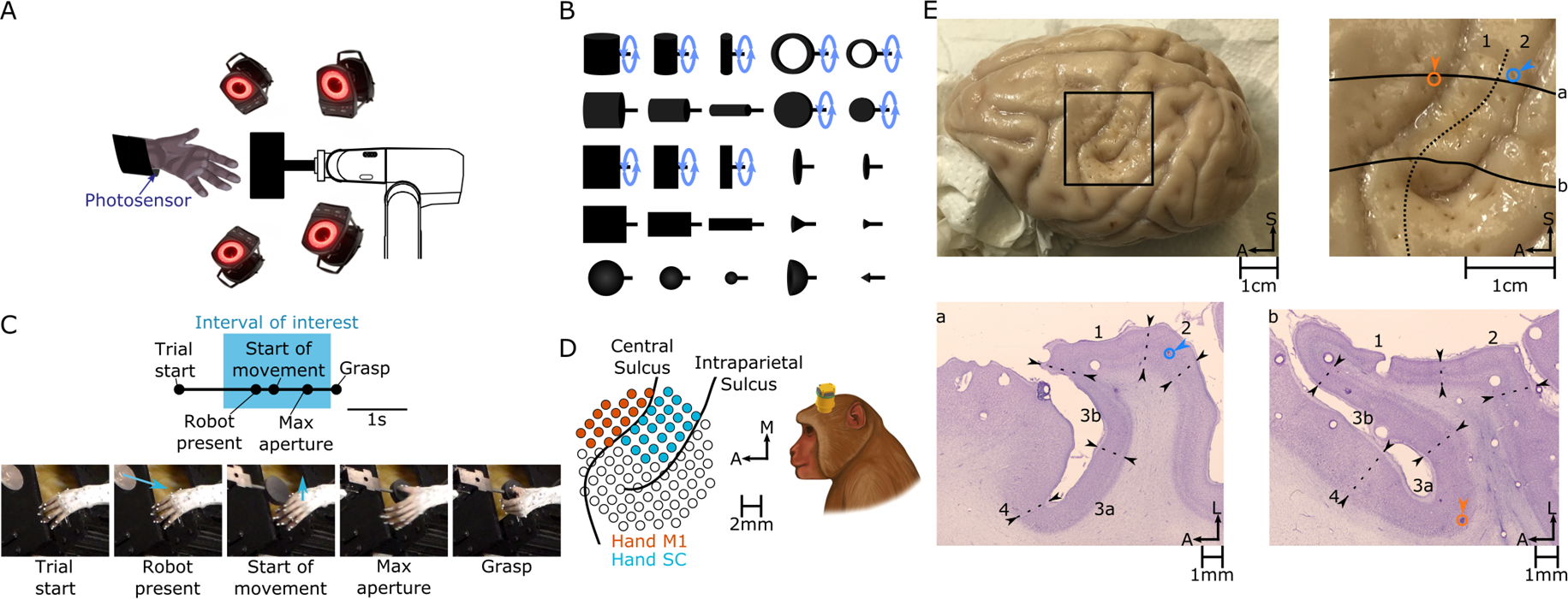
Experimental methods. **(A)** Rhesus monkeys grasp objects presented by a robotic arm. Monkeys keep their arms largely immobile, only moving the hand to grasp objects. If the monkey lifts its arm to reach for the object, a photosensor is triggered and the trial is aborted. A fourteen-camera motion tracking system tracks the kinematics of the hand. **(B)** Set of 25 shapes used in this study. Ten of these shapes (indicated by a blue circular arrow) were presented at different orientations, totaling 35 “objects”. **(C)** Task progression. The “Start of movement”, “Max aperture”, and “Grasp” events were manually scored from video. Blue arrows in photos (bottom) indicate motion of the robot (“Robot present”) or the hand (“Start of movement”). Analyses were confined to neural responses measured prior to “Grasp” to eliminate the confounding effects of object contact. **(D)** Multi-electrode arrays are used to record neuronal activity. Pictured on the left are the reconstructed locations of electrodes relative to the surface of cortex (left hemisphere) in Monkey 4. See Figure S1 for array placements in other monkeys relative to cortical surface landmarks. **(E)** Histological reconstruction of array placement. *Top left*: Chronically implanted electrode tracks were clearly visible in the perfused cortex. *Top right*: Enlarged view of the rectangular region in *top left*. Registered to this view of the cortical surface are the architectonic boundary between areas 1 and 2 (dashed line); the locations of two electrodes that putatively recorded from areas 2 (blue) and 3a (orange); and the contours of cortex along two horizontal slices pictured in *bottom* (solid lines). *Bottom*: Both horizontal slices are stained for Nissl substance, and boundaries between cortical fields are drawn on the basis of architectonic features. Specifically, the transition from area 4 to 3a coincides with a reduction in the frequency of pyramidal cells in layer 5 and an increase in the density of granular layer 4; between 3a and 3b, a further increase in the density of layer 4; between 3b and 1, an increase in the thickness of layer 4; between 1 and 2, a reduction in the density of layer 4; and at the caudal extent of area 2, a further reduction in layer 4 density. In slice **a** (*left*), the recording site of the blue electrode is confirmed to reside within the boundaries drawn for area 2. In slice **b** (*right*), the recording site of the orange electrode is confirmed to reside within area 3a.

## Results

### Kinematics (and neuronal responses) are object-dependent

To characterize the neural representation of a complex effector such as the hand requires that the space of hand postures and movements be diverse. To verify that grasping occupies a sufficiently rich space of hand kinematics, we first examined the degree to which different objects evoked different kinematics. We found that hand posture trajectories diverge long before contact is established with the object (Figure 2A). That is, the animal preshaped its hand to each object to grasp it, and this preshaping began to emerge shortly after movement onset (Figure 2B). To characterize the object-specificity of hand shape, we classified objects based on hand postures aligned to various events before object contact (Figure 2C). We found that classification performance evolved gradually over the course of the trial, was well above chance (chance performance = 2.8%) at maximum aperture, and peaked at around 60% just before grasp. This analysis confirms that animals preshape their hands over most of the duration of the trial in an object specific manner. We also examined the responses of somatosensory and motor cortical neurons and found a wide variety of response profiles (Figure S2A). The degree to which responses were object dependent varied from neuron to neuron, but sufficient information was carried in these neuronal signals to support object classification at the population level (Figure S2B,C). Thus, both hand movements and the evoked responses are diverse and object specific.

**Figure 2.**
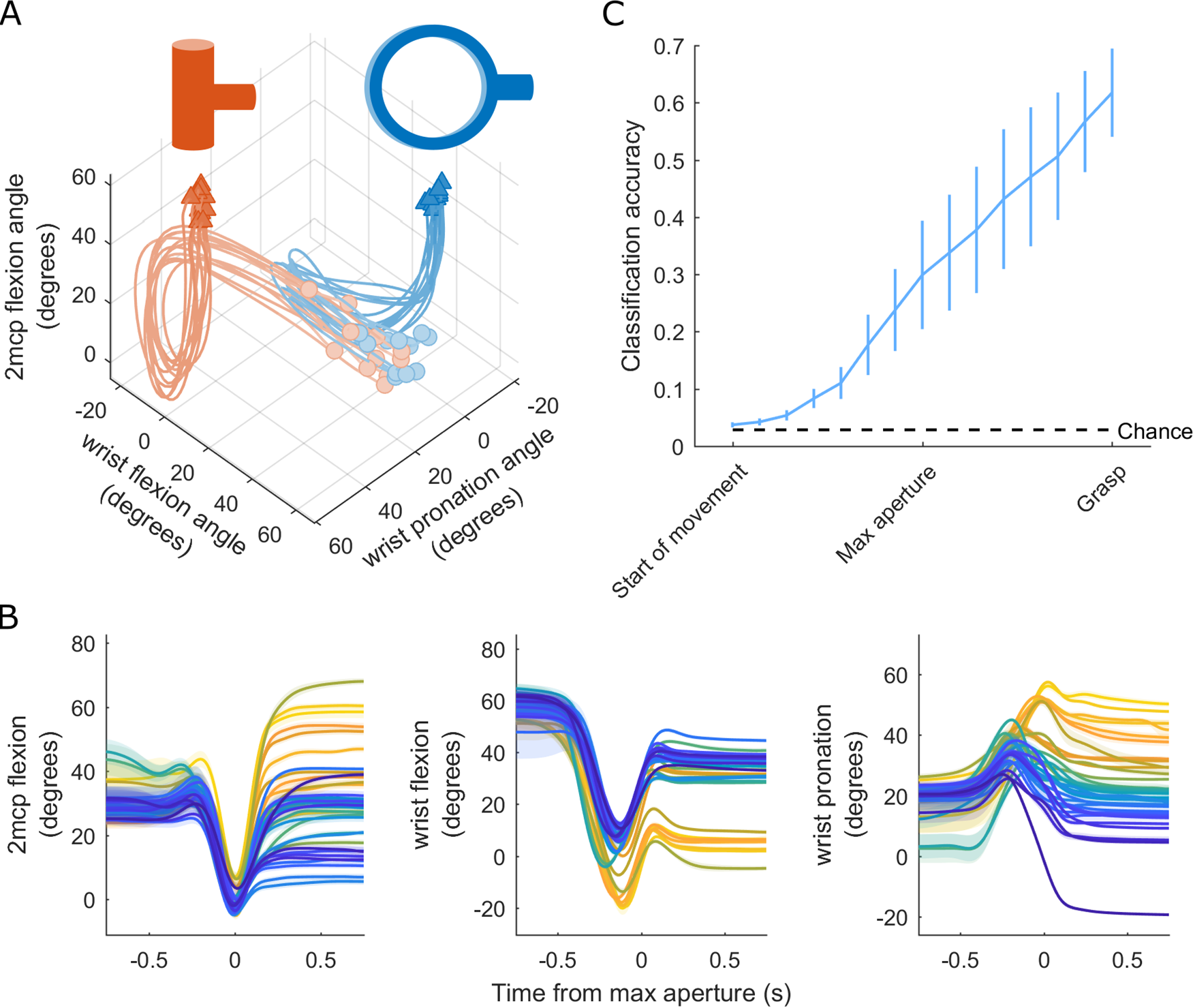
Different objects give rise to different hand pre-shaping kinematics. **(A)** Trajectories of three joints as an animal grasps two different objects over the course of a session. Each trace represents kinematics during a single trial. Faded circles indicate joint angles 750 ms prior to maximum aperture and darkened triangles indicate joint angles 750 ms after maximum aperture. **(B)** The joint angles from (A), plotted for the 35 objects, with different object-specific trajectories indicated with different colors. Each colored trace is averaged across all presentations of that object. Shading indicates ±1 S.E.M. across trials. **(C)** Time-varying object classification on the basis of the posture of the hand, assessed across all sessions and averaged across monkeys. Error bars indicate ±1 S.E.M. across monkeys. Black dashed line indicates chance performance. See Figure S2 for corresponding variety in neural responses.

### Encoding models based on kinematics can predict neuronal responses

To assess which movement features drive the responses of SC and M1 neurons, we fit a generalized linear model (GLM) to predict the time-varying firing rate of each neuron from the time-varying joint angles and their derivatives. GLMs often provided accurate predictions of neuronal responses (Figure 3A), with goodness of fits comparable to those achieved by similar models applied to proximal limb responses in motor cortex during reaching (Figure 3B, Table S1) (Hatsopoulos et al., 2007). Thus, neurons in somatosensory and motor cortex encode hand kinematics with a comparable fidelity as they do arm kinematics.

**Figure 3.**
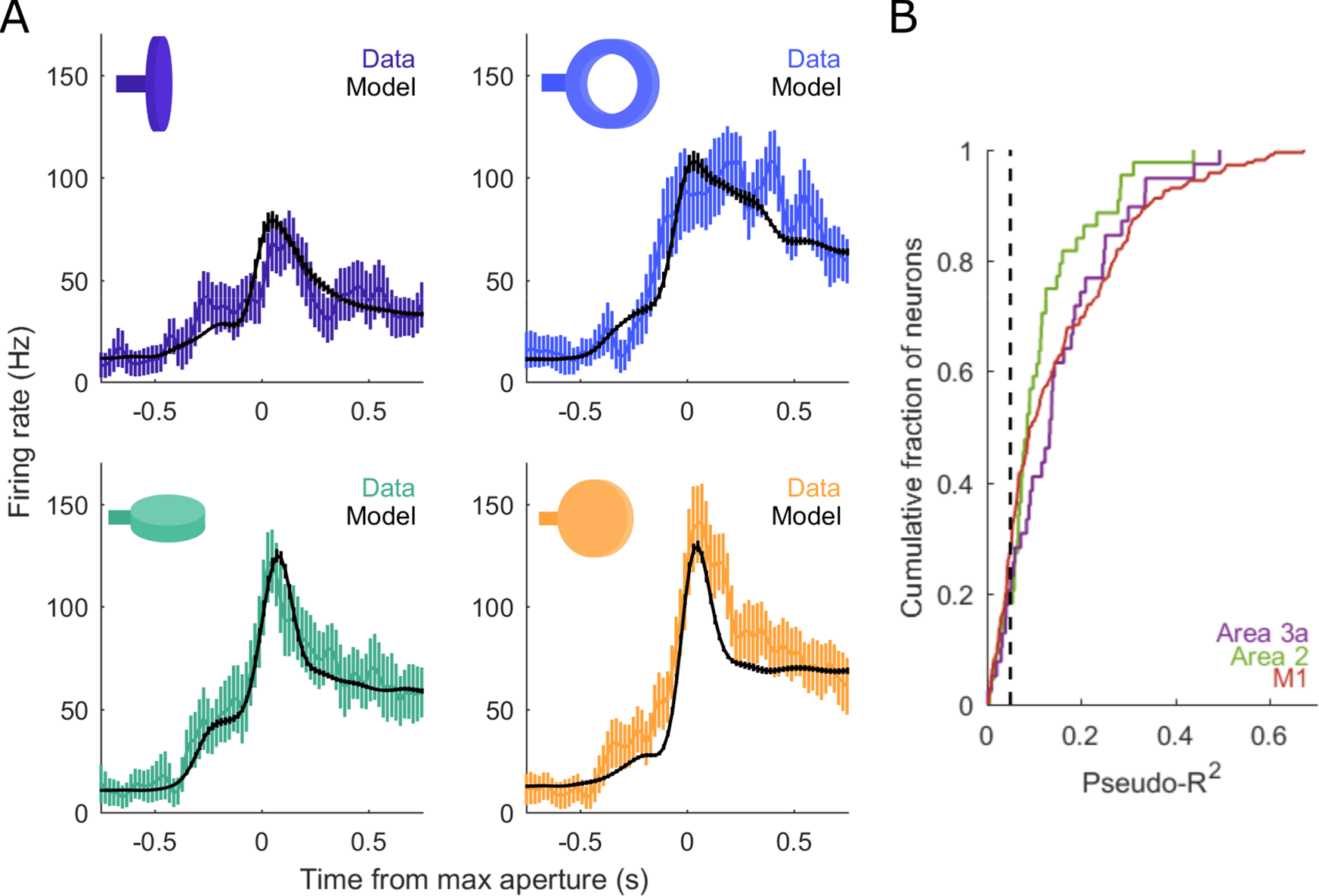
Performance of the generalized linear model (GLM). **(A)** Measured (colored) and predicted (black) peri-event time histograms (PETHs) for a single neuron in area 3a. Each plot depicts the PETH associated with a different object. The pseudo-*R^2^* of the GLM fit to this neuron is 0.34. Vertical bars indicate ±1 S.E.M. across trials. **(B)** Cumulative distribution of pseudo-*R^2^* values across neurons from each area. Neurons are pooled across sessions and across different monkeys. The black dashed line indicates a pseudo-*R^2^* cut-off of 0.05, which is used in subsequent analyses of response field (RF) structure. In area 2, the pseudo-*R^2^* values of 36 out of 44 total units (81.8%) exceed this criterion; in area 3a, 32 of 39 (82.1%); and in M1, 167 of 219 (76.3%). Importantly, the average goodness-of-fit is comparable to that reported for M1 neurons during reaching (Table S1).

### Neuronal response fields span many joints

Next, we examined which movement features drive the responses of SC and M1 neurons. First, we assessed whether individual neurons encode the state of one joint or that of multiple joints. We found that neurons encode combinations of joints, with multi-joint models accounting for roughly twice the deviance in spiking responses as did single-joint models (Figure 4A). In fact, 8 joints were required, on average, to account for 90% of the response deviance, with little difference in response field size across somatosensory and motor cortical fields (Figure 4B,C). We verified that these large response fields were not an artifact of inter-joint correlations (Figure S3A).

**Figure 4.**
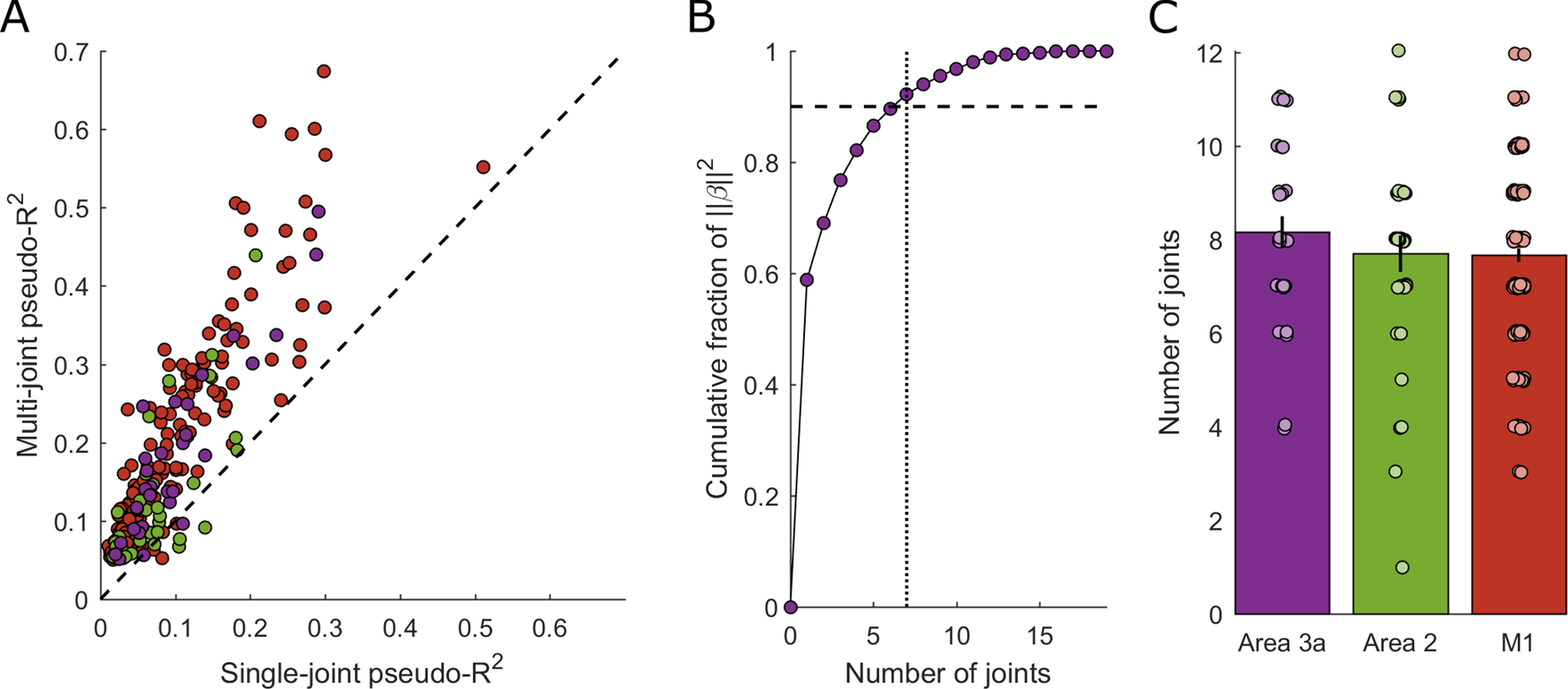
Response field size for somatosensory and motor cortical neurons. **(A)** Comparisons of each neuron’s best single-joint pseudo-*R^2^* (abscissa) against the corresponding multi-joint pseudo-*R^2^*. Multi-joint models yield considerably better predictions than do single-joint models, achieving roughly doubled levels of goodness-of-fit. Dashed line indicates the unity slope. **(B)** For each weight vector, β, defining a neuron’s response field (RF), we calculate the contribution of each joint to the squared norm of β. The minimum number of joints (dotted vertical line) required to account for 90% of that norm (dashed horizontal line) is taken to be the set of joints defining that neuron’s RF. **(C)** Average number of joints in a neuron’s RF for each area. Only neurons with pseudo-*R^2^* > 0.05 are considered. There are no differences across areas: Roughly eight joints define the typical RF from each area. Individual points denote joint counts for the response fields of individual neurons. Vertical lines indicate ±1 S.E.M. across neurons. Such distributions of joint counts are unlikely to emerge from neurons that only track or control a single joint (Figure S3A).

### Response fields span the entire hand

Next, we examined how the joints were distributed over the hand. One might expect, especially at early stages of somatosensory processing such as area 3a, that multi-joint response fields (RFs) would be confined to just one digit or two adjacent ones, controlled by a common muscle. To characterize the spatial extent of the RFs, we computed co-occurrence matrices, which show the likelihood that a given pair of joints is present in an individual neuron’s RF, conditioned upon at least one of those joints being present. We found that RFs spanned the entire hand (Figure 5A). That is, the metacarpo-phalangeal (MCP) joints of digits 1-5, the carpo-metacarpal (CMC) joints of digits 1 and 5, and the wrist all co-occurred with one another at similar rates, as opposed to forming separate clusters of co-occurring joints.

**Figure 5.**
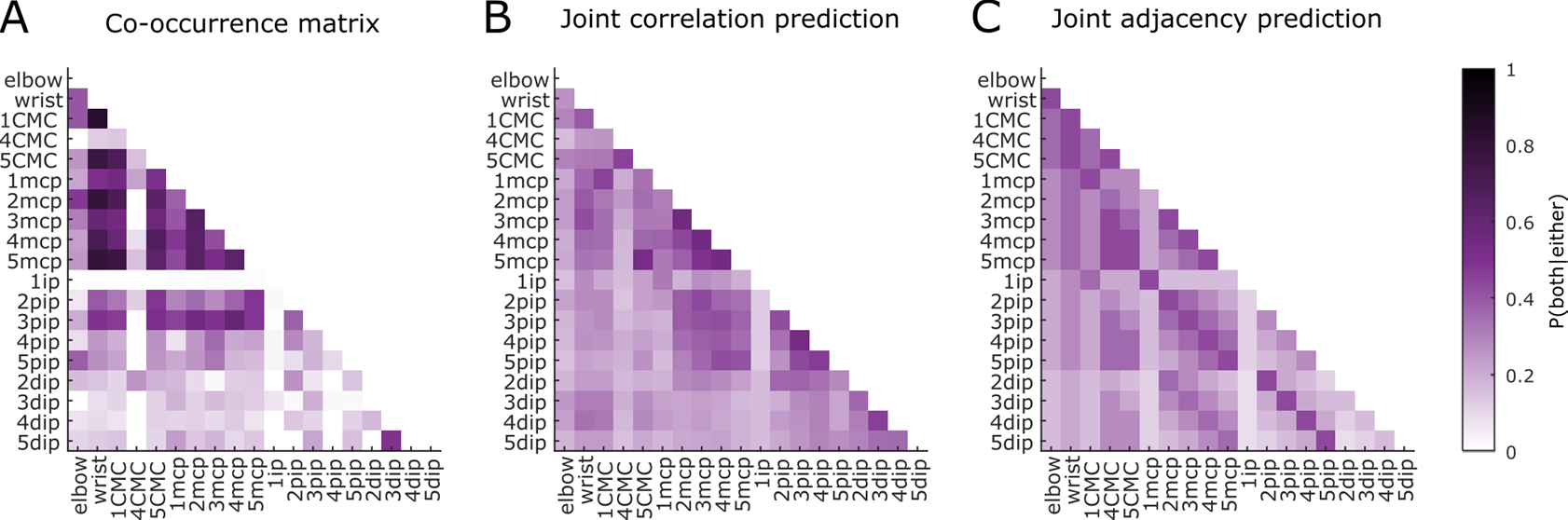
Co-occurrence matrices for area 3a. Co-occurrence is defined here as the likelihood that two joints of a pair are contained within the same RF given that at least one of them is present within that RF. **(A)** Empirical co-occurrences taken from the population of area 3a neurons with pseudo-*R^2^* > 0.05. **(B)** Co-occurrences predicted from correlations between joints. This model fails to explain a large fraction of the deviance in the co-occurrence data (*R^2^* = 0.239). **(C)** Co-occurrences predicted from joint proximity. Predictions are based on minimum path length between joints in the undirected, unweighted “adjacency” graph whose edges are defined by pairs of joints that share at least one bone between them, or those that comprise adjacent metacarpal bones (bones defining joints are provided in Table S2; details governing “adjacency” are explained in the Methods). Only area 3a is shown in this figure; a similar analysis was performed for area 2 and M1 as well, all yielding similar conclusions (Figure S3B): Namely, that co-occurrence of pairs of joints in neuronal RFs is not adequately predicted from just the correlations between those joints or the proximity of those joints (*R^2^* = 0.146).

We then examined whether patterns of co-occurrence in neuronal RFs trivially reflected kinematic correlations among joints (Figure 5B) or the anatomical proximity of those joints (Figure 5C). We found that neither model could account for the patterns of co-occurrence observed in area 3a (Figure 5), area 2, or M1 (Figure S3B).

To further test whether natural correlations among joints might be shaping the large RFs of proprioceptive neurons, we computed the principal components (PCs) of the kinematics – which reflect correlated patterns of joint postures and are sometimes inferred to be canonical “synergies” or “movement primitives” from which all movements arise (Santello et al., 1998). We regressed these, rather than the joint angles themselves, onto the neural response. If such a coordinate frame were preferentially encoded, one might expect individual PCs to predict firing rates better than individual joints, or for multi-PC models to be more parsimonious — i.e., require fewer parameters — than multi-joint models. We found that individual PCs are not preferentially encoded over individual joint angles in proprioceptive neurons; rather, single-joint models actually slightly outperformed their single-PC counterparts (paired-samples t-test, *t*(82)=2.946, *p*=4.19e-03) (Figure 6A). Moreover, PC space did not provide a more parsimonious model of neuronal firing rates than did the full joint space (*t*(226)=0.361, *p*=0.718) (Figure 6B). This was true in all cortical fields, and results from M1 replicate previous findings with a different manual task (Kirsch et al., 2014; Mollazadeh et al., 2014). Sensorimotor cortical fields do not seem to preferentially represent kinematics in a principal component coordinate frame relative to a joint coordinate frame.

**Figure 6.**
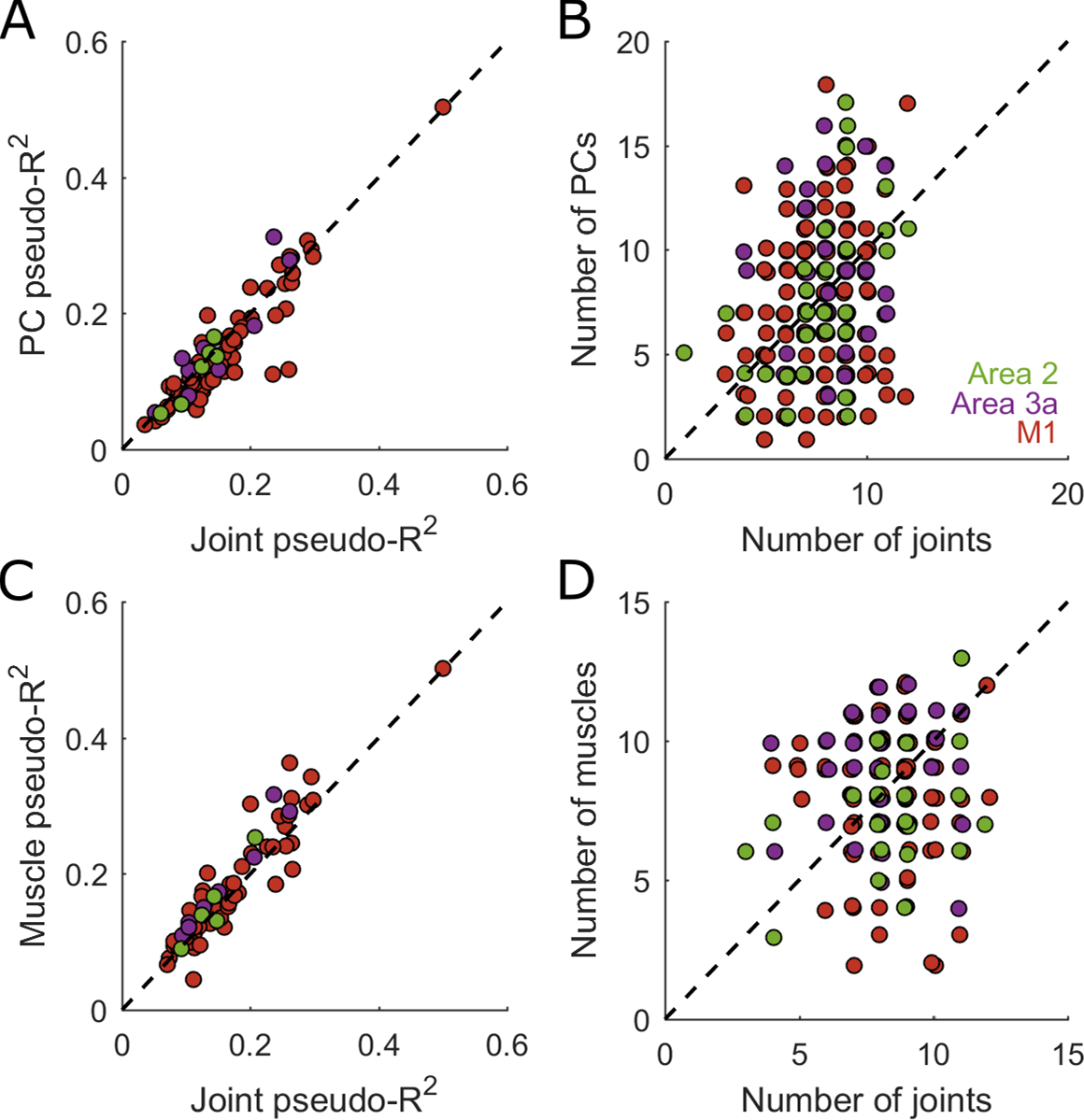
Lack of parsimonious RF description across kinematic coordinate frames. **(A)** The best single-joint GLM fit to the most strongly-modulated neurons does no worse than the best single-PC GLM in terms of pseudo-*R^2^* of predicted firing rates. Only neurons with pseudo-*R^2^* > 0.20 with the multi-joint GLM are considered to avoid comparing potentially unstable models. **(B)** Roughly the same number of PCs and joints are required to achieve similar levels of performance with multi-PC or multi-joint models. Here, all models with pseudo-*R^2^* > 0.05 are considered because multiple predictors are allowed. **(C)** Comparison of the best single-joint against the best single-muscle GLM. Only neurons with pseudo-*R^2^* > 0.20 are included. **(D)** Comparison of the number of joints against number of muscles in multi-predictor models. Models with pseudo-*R^2^* > 0.05 are included.

Finally, as proprioceptive signals are thought to emanate primarily from muscle and tendon-associated receptors (spindles and Golgi tendon organs), we examined whether the musculotendon lengths of extrinsic hand muscles were preferentially encoded in cortical responses over joint kinematics. We found that individual muscles accounted for only slightly (though significantly) more deviance in firing rate than did individual joints (paired-samples t-test, *t*(69)=3.220 *p*=1.96e-03) (Figure 6C). However, multi-muscle models of spiking activity were no more parsimonious — i.e., required no fewer parameters to explain the neural response — than were multi-joint models (*t*(162)=0.256, *p*=0.798) (Figure 6D). We conclude that the RFs of proprioceptive neurons span complex combinations of joint angles, kinematic principal components, and musculotendon lengths, with no clear “labeled line” code preferentially emerging in any one kinematic reference frame.

### Neurons encode hand postures rather than movements

Individual neurons in SC and M1 respond much more strongly when the arm is moving than when it is not (London and Miller, 2013; Moran and Schwartz, 1999; Paninski et al., 2004; Reina et al., 2001; Wang et al., 2007; Weber et al., 2011). In light of this, we assessed whether hand representations are also dominated by movements over postures (Figure 7A). To this end, we analyzed the partial pseudo-*R*^2^ values for postural and movement encoding models of each neuron’s responses (Movshon and Newsome, 1996), or the fraction of unique deviance explained (FUDE) by each of these models, and found postures to dominate movements (Figure 7B). Our results therefore suggest that proprioceptive neurons preferentially track postures, not movements, of the hand, unlike their counterparts that encode arm movements (Figure S4) (Hatsopoulos et al., 2007). Moreover, and surprisingly, M1 neurons also exhibit a strong preference for hand postures during grasp, in stark contrast to the extensively documented movement preference during reaching and to earlier reports suggesting movement preference during grasp (Saleh et al., 2010) (see Figure S5 for further validation of this result).

**Figure 7.**
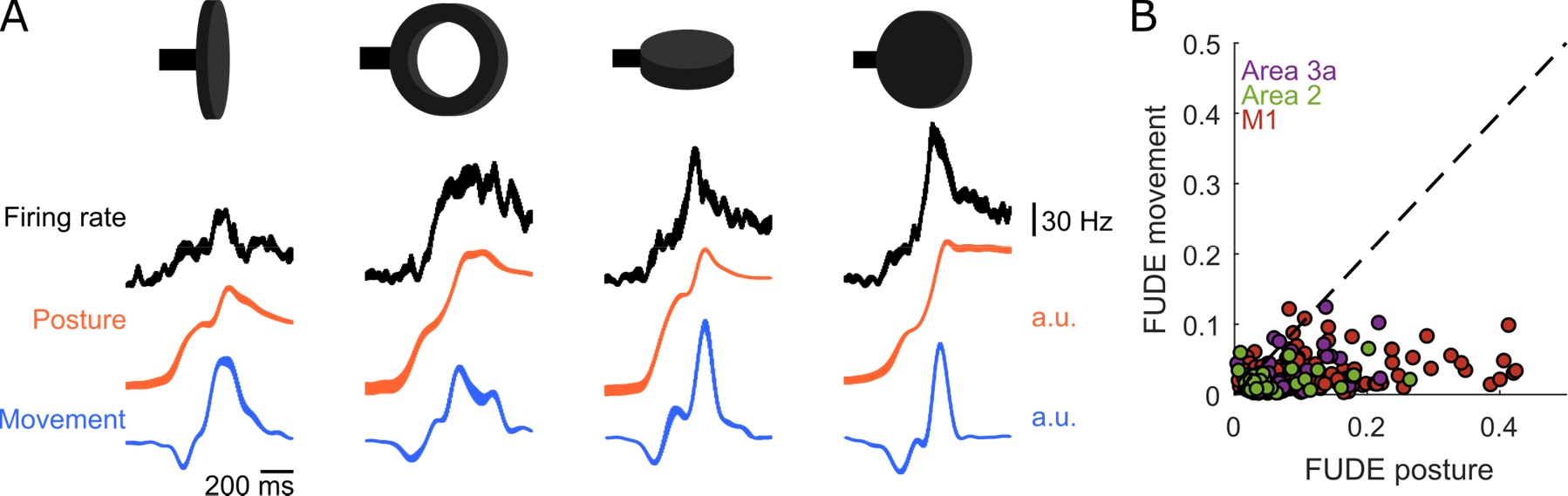
Preferential encoding of joint postures in somatosensory and motor cortex. **(A)** Peri-event time histograms from Figure 3A (*top row*), shown with hand postures (*middle row*) or movements (*bottom row*) along the respective dimension most aligned with the neuron’s firing rate. Best dimensions for postures and movements are fit using separate generalized linear models (GLMs), one using only postural predictors, and one using only movement ones. Being along separate axes, the posture and movement traces are not directly derived from one another. Qualitatively, both the posture and velocity axes account for the second, larger response transient occurring at maximum aperture, but the postural axis better captures the first response transient prior to maximum aperture and sustained activity following maximum aperture. **(B)** The fraction of unique deviance explained (FUDE) by postural kinematics (posture) and movement kinematics (movement). Each point represents a single neuron. That the points all fall well below the unity line (dashed line) suggests that postures, rather than movements, are preferentially encoded by these neurons. Note that this does not hold for M1 during reaching movements (Figure S4). See Figure S5 for further verification of the posture-encoding result, particularly in light of previous reports on preferential coding along movement axes in M1 during grasp (Saleh et al., 2010).

## Discussion

### Large postural response fields in somatosensory and motor cortex

Proprioceptive and motor representations of the hand preferentially encode postures over movement during grasping, in contrast to their proximal limb counterparts during reaching (London and Miller, 2013; Moran and Schwartz, 1999; Paninski et al., 2004; Reina et al., 2001; Wang et al., 2007; Weber et al., 2011). These differences might reflect different constraints on movement imposed by the different inertial and biomechanical properties of the arm and hand (Gribble and Scott, 2002; Kalaska et al., 1989; Prud’homme and Kalaska, 1994; Sergio et al., 2005). Indeed, the mass of the entire arm is large whereas that of the digits is negligible. The two effectors would thus require different muscle recruitment profiles to achieve similar kinematics. These biomechanical factors may thus underlie the observed differences in neuronal firing profiles for reaching and grasping. However, proprioceptive afferents with response fields on the hand are more sensitive to movements than postures (Dimitriou and Edin, 2008a, 2008b). The difference in postural sensitivity observed in cortical proprioceptive representations of the arm and hand is thus not inherited from the periphery, as would be predicted if it were an artifact of biomechanics. These results imply that proprioceptive representations of the hand are transformed from movement-dominated to posture-dominated as one ascends the neuraxis. Whether this transformation is first achieved in cortex or inherited from upstream structures – the cuneate nucleus or somatosensory thalamus – remains to be determined.

Another property of proprioceptive neurons in SC is that their response fields include several joints spanning the entire hand, as do their counterparts in M1 (Costanzo and Gardner, 1981; Saleh et al., 2010, Schieber, 1996, 2001; Schieber and Hibbard, 1993). The large response fields in area 3a stand in stark contrast to the small tactile receptive fields in somatosensory cortex (Pons et al., 1985), but consistent with previous reports of larger RFs in area 3a relative to tactile area 3b (Krubitzer et al., 2004). However, we find that a much larger proportion of proprioceptive neurons have large multi-digit RFs than previously reported (Costanzo and Gardner, 1981; Krubitzer et al., 2004), a difference that could be attributed to our analysis of responses during actively-generated natural grasping movements, rather than movements imposed on the joints.

Differences in proprioceptive and cutaneous response field sizes may reflect differences in the function of these two sources of somatosensory input. Indeed, cutaneous signals convey information about local shape features at the points of contact between hand and object such as edge orientation (Bensmaia et al., 2008) and curvature (Yau et al., 2013). Smaller receptive fields lead to more acute tactile spatial representations. Proprioceptive signals convey information about the configuration of the hand, which by definition requires integrating information across the entire hand.

Ultimately, these two streams of information – tactile and proprioceptive – must be integrated. Indeed, local features of the object at each point of contact must be interpreted in terms of the relative positions of the contact points in three dimensional space, to culminate in stereognosis, a three dimensional percept of the object (Delhaye et al., 2018). That individual neurons signal the postures of joints distributed over the entire hand is consistent with a view that the hand representation in SC and M1 emphasizes the configurations of the digits relative to one another, a representation that is ideally suited to support stereognosis.

### Alternate coordinate frames fail to account succinctly for large response fields

We investigated whether the large proprioceptive and motor response fields could be more parsimoniously accounted for in a different coordinate frame. One hypothesis is that multi-digit response fields are mediated by one or two extrinsic muscles of the hand. For example, the extensor digitorum communis inserts onto all digits and even influences wrist movements. We found, however, that models based on musculotendon lengths only modestly increased the accuracy of single-predictor models and are not more parsimonious.

Another possibility is that proprioceptive neurons encode correlated combinations of joints. Indeed, according to some variants of the synergy hypothesis, neural control of the limb is restricted to a lower dimensional manifold than that afforded by its biomechanics to render its control simpler and more robust to motor noise (Flash and Hochner, 2005; Kutch and Valero-Cuevas, 2012; Tresch and Jarc, 2009). This putative manifold is revealed through standard dimensionality reduction approaches such as principal component analysis. The efficient coding hypothesis (Barlow, 1961) would prescribe that the associated sensory system would also reflect this lower dimensional manifold so that redundancies in the hand postures would not be redundantly represented in sensory cortex. Such a representation would also be supported by Hebbian synaptic learning, which is known to give rise to neuronal representations that reflect correlated patterns of inputs (Friston et al., 1993; Miller and MacKay, 1994; Oja, 1982, 1992; Pehlevan et al., 2015) — in this case, joint kinematics. Against these predictions, however, we found that models based on principal components of the kinematics – which collapse kinematic redundancy into a basis set of non-redundant signals – were not more accurate or parsimonious than models based on raw joint angles, consistent with previously findings in M1 (Kirsch et al., 2014; Mollazadeh et al., 2014).

Additionally, neuronal representations may reflect limb dynamics (i.e., the underlying forces) rather than kinematics. For example, limb forces and muscle activations may be better encoded in M1 than are kinematics (Evarts, 1968; Morrow et al., 2007; Sergio et al., 2005). Additionally, limb forces may be necessary to account for the responses of a subset of neurons in somatosensory cortex (Prud’homme and Kalaska, 1994). Because we cannot accurately reconstruct muscle activations and dynamics from kinematics, we cannot test this hypothesis directly. However, the activity of single neurons in motor cortex has been shown to drive suppression and facilitation of several muscles at once (Buys et al., 1986; Griffin et al., 2015; Hudson et al., 2017) so there is no reason to believe that an encoding model based on hand dynamics or muscle activations would be more parsimonious than one based on kinematics.

More parsimonious models are likely to arise in a coordinate frame that captures behaviorally relevant features, such as hand aperture (Jeannerod, 2009; Jones and Lederman, 2006) or grip type (such as “power” and “precision” (Napier, 1956)) or one that explicitly represents the relative positions of the fingertips (Biggs et al., 1999). Similar transformations of coordinate frames have revealed parsimonious descriptions of otherwise complex neural response properties in macaque face patches (Chang and Tsao, 2017). A major challenge moving forward is to understand how hand postures are encoded along the somatosensory neuraxis and how these postural representations interact with cutaneous representations of object contacts to give rise to stereognosis (Delhaye et al., 2018; Hsiao, 2008).

### Proprioceptive and motor representations of the hand are similar

Neurons in SC and M1 carry remarkably similar representations of the hand, as has been found for the proximal limb (London and Miller, 2013). Hand response fields are of approximately equal size and the degree of posture-preference is approximately equivalent in neurons confirmed by histology (Figure 1E) to reside in Brodmann’s areas 3a, 2, and 4 (M1). Note that one canonical difference between motor and sensory responses – that the former precede and the latter follow movement – is difficult to probe with natural movements given autocorrelations that stretch over long time scales, particularly in the hand postural kinematics preferentially tracked by somatosensory and motor cortices during grasp (Figure S5E).

The similarity between proprioceptive and motor representations may reflect their tight interplay in the neural control of movement. Indeed, somatosensory cortex is essential for the execution of skilled movement (Brochier et al., 1999b; Hikosaka et al., 1985; Jeannerod et al., 1984) and area 3a is known to send and receive projections directly to and from motor cortex in primates (Huerta and Pons, 1990; Huffman and Krubitzer, 2001). The bidirectional communication between somatosensory and motor cortices may be facilitated by a common representational scheme for the hand.

### Conclusions

Proprioceptive representations of the hand encode time-varying joint postures distributed over the entire hand. This neural representation stands in contrast with its proximal limb counterpart, which preferentially encodes movement, a difference that does not seem to be inherited from the periphery. Proprioceptive response fields in SC, similar to their counterparts in M1, cannot be straightforwardly explained by patterns in the hand movements or by the anatomy of the hand. The postural representation of the hand in sensorimotor cortex is consistent with the primacy of tracking hand shape and for a role of these representations in stereognosis.

## Methods

### Animals and surgery

We recorded neural data from four male Rhesus macaques (Macaca mulatta) ranging in age from 6 to 15 years and weighing between 8 and 11 kg. All animal procedures were performed in accordance with the rules and regulations of the University of Chicago Animal Care and Use Committee (IACUC). Monkeys received care from a full-time husbandry staff, who maintained a 12hr/12hr light/dark cycle, cleaned the animals’ living spaces once a week, and provided the animals with ample food and enrichment. In addition, a full-time veterinary staff monitored the animals’ health. Animals were water-restricted according to a protocol requiring monitoring their weights daily and ensuring an absolute daily minimum water consumption of 10 cc/kg.

Monkeys underwent a magnetic resonance (MRI) scan to identify anatomic landmarks and stereotaxic coordinates in preparation for array implantation. Each monkey was then implanted with a head post fixed to the skull with bone screws. Monkey 1 was implanted with two Utah electrode arrays (UEAs, Blackrock Microsystems, Inc., Salt Lake City, UT), one in primary motor cortex, the other in somatosensory cortex and four floating microelectrode arrays (FMAs, Microprobes for Life Science, Gaithersburg, MD), two in the anterior and two in the posterior bank of the central sulcus (Figure S1A). Monkeys 2 through 4 were implanted with semichronic Microdrive electrode arrays (SC96, Gray Matter Research, Bozeman, MT), each spanning large swaths of primary motor and somatosensory cortex and comprising individually depth-adjustable electrodes (Figure S1B-D) (Dotson et al., 2017). All procedures were performed under aseptic conditions and under anesthesia induced with ketamine HCl (20 mg/kg, IM) and maintained with isoflurane (10-25 mg/kg per hour, inhaled).

### Behavioral task and recording methods

Animals were trained to perform a grasping task. On each trial, one of 25 objects was brought to the animal’s stationary hand by an industrial robot (MELFA RV-1A, Mitsubishi Electric, Tokyo, Japan) (Figure 1A). As the object approached, the animal preshaped its hand to grasp it. Some of the objects were presented at different orientations, requiring a different grasping strategy, so the different orientations of the same object will be referred to as different objects (Figure 1B). Each object (of the 35 “objects”) was presented eight to eleven times in a given session. A set of 31 reflective markers was placed on bony landmarks straddling the joints of the hand and forearm (Figure S1E,F) and a 14-camera optical tracking system (MX T-Series, VICON, Los Angeles, CA) (Figure 1A) tracked their time-varying three-dimensional positions at a sampling rate of 250 Hz (Monkey 1) or 100 Hz (Monkeys 2-4).

Different trial epochs could be divided based on five events (Figure 1C): Trial start, when the cameras began to record kinematics; robot present, when the object began to move toward the monkey’s hand; start of movement, the time at which the hand began to move about the wrist joint; maximum aperture, a critical component of hand pre-shaping (Jeannerod, 2009; Jones and Lederman, 2006), the time at which the digits were maximally separated; and grasp, when object contact was finally established. Only neural data during pre-grasp epochs, extending from 750 ms prior to start of movement through 10 ms prior to grasp, were analyzed. The timing of start of movement, maximum aperture, and grasp events were inferred on the basis of the recorded kinematics. A subset of trials from each session were manually scored for each of these three events. On the basis of these training data, joint angular kinematic trajectories spanning 200 ms before and after each frame were used as features to train a multi-class linear discriminant classifier to discriminate among these four classes: all three events of interest and “no event”. Log likelihood ratio was used to determine which “start of movement”, “maximum aperture”, and “grasp” times were most probable relative to “no event”. Events were sequentially labeled for each trial to enforce the constraint that start of movement precedes maximum aperture, and maximum aperture precedes grasp.

After grasp, monkeys were required to maintain contact with the object for an interval lasting between one and three seconds (randomly drawn on each trial), after which the robot would retract. If the monkey maintained enough grip force to disengage the magnetic coupling between robot and object during the retraction of the robot, a water reward was administered.

Because we wished to investigate coding of hand movements, we sought to eliminate movements of the proximal arm associated with reaching, which can overlap substantially with grasp representations, especially in motor representations of the limb (Donoghue et al., 1992; Mckiernan et al., 1998; Park et al., 2001; Saleh et al.; Takahashi et al., 2017). However, the use of restraints to hold the arm in place would introduce cutaneous inputs and isometric forces exerted against those restraints, which might affect neural responses but not be reflected in the measured kinematics. To minimize these confounds while still isolating grasping movements, we trained monkeys to volitionally hold their arms stationary while grasping objects. This was achieved by placing a photosensor under each arm (Figure 1A) and only rewarding the monkeys when they performed the task without moving their arms off the sensors.

In addition to kinematic data, neural data were recorded from neurons in somatosensory and primary motor cortex using multi-electrode arrays. Somatosensory cortex (SC) comprises four cortical fields, each containing its own body map (Kaas et al., 1979; Pons et al., 1985): Areas 3a, 3b, 1 and 2. Measurements were focused on areas 3a and 2, which are known to contain neurons with proprioceptive responses (Jones and Porter, 1980; Kaas, 1983; Pons et al., 1985). In Monkey 1, UEAs targeted cortical fields on the post- and pre-central gyri and FMAs targeted cortical fields located deep in the banks of the central sulcus, namely caudal M1 (anterior) and area 3a (posterior)(Figure S1A). In Monkeys 2-4, each SC96 array impinged on all relevant areas of SC and M1 given the wide span of this implant and the fact that its comprises depth-adjustable electrodes (Figure 1D) (Figure S1B-D). Histological reconstructions, obtained for one monkey (Monkey 4), confirmed the location of proprioceptive neurons in Brodmann’s areas 3a or 2 (Figure 1E).

### Data Processing

We used marker positions during grasp to reconstruct joint angles and musculotendon lengths. Inverse kinematics were calculated using labelled marker kinematics and a musculoskeletal model of the human arm (https://simtk.org/projects/ulb_project) (Anderson & Pandy, 2001; Anderson & Pandy, 1999; de Leva, 1996; S.L. Delp et al., 1990; Dempster & Gaughran, 1967; Holzbaur, Murray, & Delp, 2005; Yamaguchi & Zajac, 1989) implemented in Opensim (https://simtk.org/frs/index.php?group_id=91) (Delp et al., 2007) and scaled down to the size of a monkey arm. Inverse kinematics returned estimates of the time-varying joint angular coordinates—22 in Monkey 1 and 30 in Monkeys 2-4, spanning all degrees of freedom across 13 joints in Monkey 1, and 19 joints in Monkeys 2-4. Kinematic reconstruction of the extrinsic muscles of the hand was also possible for Monkeys 2-4. Details of the joints and muscles we reconstructed are given in Table S2.

Inverse kinematic data were filtered first using a moving median filter (MATLAB movmedian) over a centered 83 ms window to remove outliers and sudden jumps in the kinematic data. The output of the moving median filter was then filtered using a 4^th^ order low-pass Butterworth filter with a 6 Hz cutoff frequency (MATLAB butter and filtfilt). Joint angular velocities were then calculated from these filtered kinematics (MATLAB diff).

Spikes in the neural data were detected by first identifying manually-set threshold crossings in the raw voltage trace, sampled at 30 kHz and digitally high-pass filtered with a cutoff frequency of 200 Hz. Offline spike sorting (Offline Sorter, Plexon, Dallas, TX) was then used to isolate individual units from a trace if more than one action potential waveform was identified and to remove non-spike threshold crossings.

### Functional hand mapping

As our interest was primarily in hand proprioceptive responses in somatosensory cortex, we functionally mapped somatosensory cortical units for proprioceptive responses. To this end, we first manually palpated the arm, hand, face, trunk, and legs and only accepted neurons responding selectively to palpations of the upper limb. We then applied light cutaneous stimulation by brushing the hand and arm tangentially with a cotton swab, and subsequently manipulated the joints of the hand and wrist and palpated the tendons and bodies of the forearm musculature. In some cases, tactile stimulation was applied to the tactile receptive field of the neuron while the hand was shaped in different postures or the wrist adopted positions at various degrees of flexion, extension, pronation, or supination. We recorded from somatosensory neurons that could be driven reliably by joint manipulations or forearm palpations.

### Histology

At the conclusion of electrophysiological recordings in Monkey 4, we processed the cortex for histology to confirm the locations of electrodes relative to cytoarchitectonic cortical fields. Electrolytic lesions (10 µA monophasic pulses at 300 Hz for 10 s) were placed at strategic locations across the array to help locate selected electrode tracks. The animal was then euthanized (60 mg/kg pentobarbital sodium) and transcardially perfused with 0.9% saline followed by 3% paraformaldehyde. At the end of perfusion, the brain was removed from the cranium, blocked, and left to soak overnight in 30% sucrose phosphate buffer. Next, a cryostat was used to take transverse sections of blocked cortex at a thickness of 60 µm per slice. One of every 12 slices was stained for Nissl and VGlut2, and one of every 6 for Cytochrome Oxidase (CO), to aid in identifying boundaries between cortical areas using cell body morphology and density.

A three-dimensional reconstruction of histological sections, borders of cortical fields, and electrode tracks was achieved by registering transverse slices and histological sections in ImageJ (StackReg: http://bigwww.epfl.ch/thevenaz/stackreg/ then marking each electrode or cortical field border as a region of interest (ROI). Example Nissl stains from two slices show electrode tracks in Brodmann’s areas 3a and 2, confirming the location of the recorded neurons initially estimated based on anatomical landmarks and response properties (Figure 1E).

### Data Analysis

#### Computing trial-averaged kinematics and firing rates

Kinematics and spike counts were collected across multiple presentations (~10) of each object and averaged to obtain an estimate of the time-varying hand shape (e.g., Figure 2B) or time-varying firing rates (e.g., Figure S2A) associated with each object. Prior to averaging across trials, each time-varying joint angular or spike count trace was aligned to maximum aperture and smoothed with a Gaussian kernel (σ = 20 ms). Alignment to maximum aperture yielded the largest object-dependence in the kinematics and spiking as gauged by the classification analysis (see below). Smoothed, trial-averaged kinematics and firing rates are only used for visualizing the activity of single neurons. All analyses are based on single-trial neuronal responses or kinematics.

### Kinematic object classification

To assess the degree to which the animals preshaped their hands in an object-specific way before grasp, we computed the accuracy with which classifiers could identify an object on the basis of joint angular postures at various epochs before object contact. First, we defined 15 events, 8 evenly spaced between the start of movement and max aperture, and another 7 from max aperture to grasp, and extracted the instantaneous multi-joint posture at that event for each trial. Then, we fit multiclass linear discriminant classifiers (MATLAB fitcdiscr) using multiple trials (~350) across 35 objects, and assessed classification accuracy using leave-one-trial-out cross-validation. A separate classifier was used for each epoch.

Having verified that individual monkeys used consistent grasping strategies for each object across sessions, we pooled trials across sessions from each monkey to obtain time-varying classification accuracy. We did not pool across monkeys as different grasping strategies are used by different animals. We then averaged time-varying classification accuracy across monkeys (e.g., Figure 2C) to get an overall measure of the object specificity of hand postures during pre-shaping.

### Neural object classification

Next, we wished to assess the degree to which the neuronal responses in sensorimotor cortex were object-specific before grasp. To this end, we classified objects on the basis of time-varying neural responses using similar methods as those described above for kinematics-based classification (Figure S2). The difference is that, rather than being able to concatenate trials across different sessions containing the same joints, we had to pool sessions in a manner that allows us to combine disparate sets of neurons recorded across sessions. To this end, we assumed conditional independence of each neuron (Schneidman et al., 2003), which allowed us to fit linear discriminant classifiers on a neuron-by-neuron basis and obtain an estimate of the population decoding performance by maximizing the summed log likelihood across all single-neuron classifiers. Again, we pooled across sessions within each monkey under the assumption that similar grasping strategies, and therefore similar neural population responses, were adopted by a monkey across different sessions, but did not pool across different monkeys. Additionally, rather than using instantaneous firing rate for classification, we used spike counts over a 500-ms time window preceding each event.

### Generalized linear model (GLM)

One of the main goals of the present study was to establish which aspects of hand postures and movements drove the responses of individual sensorimotor neurons. To this end, we used Generalized Linear Models (GLMs) to predict the neuronal responses over the epochs of interest (c.f. Figure 1C), from 700 ms prior to start of movement through 10 ms prior to object contact. Kinematics were aligned with these neural data at different latencies, spanning 250-ms leads of through 250-ms lags, in an attempt to find the latency that maximized the goodness-of-fit of each neuron’s GLM. In particular, we tested latencies of 0, ±10, ±20, ±30, ±50, ±90, ±150, and ±250 ms.

Note that each of 15 single-lag models fit to each neuron imposed the same uniform latency across all kinematic predictors. We did, however, assess the extent to which kinematic trajectories spanning multiple latencies influenced goodness-of-fit (Figure S5A) and found a miniscule, albeit significant, improvement with multiple lags, even after cross-validation (Figure S5). Because the effect of using multiple lags was small when expressed in terms of pseudo-*R^2^* (see below), and because single-lag models incorporate fewer parameters and are thus more readily interpretable, we used single-lag models to determine response field sizes and preferences for postural or movement kinematics.

We fit a number of different GLMs to the responses of each neuron: postural, movement, and combined. In Monkeys 2 through 4, a total of 30 predictors were used for “postural” GLMs: one for the angle about each joint degree of freedom (DOF). Similarly, 30 predictors were used for “movement” GLMs: one for the angular velocity about each joint DOF. Finally, the “combined” model used a total of 60 predictors, using both the joint angular and angular velocity predictors of the “posture” and “movement” models. In Monkey 1, 22 predictors were used in “posture” and “movement” models, and 44 in the “combined” model, as some joint degrees of freedom were not reconstructed for this monkey (Table S2).

Neural data and kinematics were down-sampled to 50 Hz (20 ms bins) prior to running the GLMs. GLMs were fit using a Poisson residual model and a softplus inverse link function and implemented in MATLAB using the nonlinear input model (NIM) (http://neurotheory.umd.edu/nimcode) (Mcfarland et al., 2013).

We use LASSO regularization to limit the number of predictors in the models. This approach introduces a hyperparameter, λ, that penalizes the L1-norm of the GLM predictor vector, β. Values of λ and β are fit using 60-20-20 cross-validation. For each λ tested, we fit the optimal β to a training set of 60% of our samples within a given session, chosen at random but kept consistent across different λ. We then estimated the log likelihood of each model on a validation set comprising 20% of our samples, chosen at random from the remaining 40% of samples not used to train each model. The λ and corresponding β that maximized the log likelihood of the validation set were then selected, and the log likelihood of the test set, comprising the final 20% of our samples, is reported.

For ease of interpretation, we then convert these log likelihoods into pseudo-*R^2^* values,

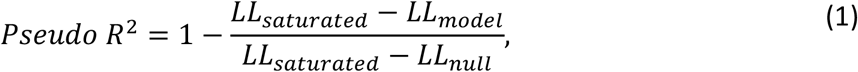

where *LL_model_* is the log likelihood of the test data given the optimal λ and β; *LL_saturated_* is the log likelihood that would be computed if spike count predictions precisely matched spike counts at each 20ms bin; and *LL_null_* is the log likelihood of the test data given a constant, non-time-varying mean firing rate. For typical linear regression, which assumes a Gaussian residual model, this pseudo-*R^2^* is precisely equivalent to the coefficient of determination, *R^2^*. For a model with Poisson residuals, it quantifies the fraction of deviance, rather than variance, explained by the GLM.

### Interpreting encoding models: size and extent of joint RFs

We computed a summary statistic to determine the number of joints in a typical neuron’s response field. We also computed partial pseudo-*R^2^* values of posture-and movement-based GLMs to determine their relative contributions of the model predictions. Summary statistics and partial pseudo-*R^2^* comparisons were computed only for the subset of neurons with a maximum cross-validated pseudo-*R*^2^ value of at least 0.05. This criterion was chosen because it is close to the *R^2^* significance criterion of regression models fit to trial-averaged data (roughly *R^2^*>0.08, c.f. Paninski et al., 2004) and is more conservative than one determined based strictly on statistical significance. In practice, the vast majority of neurons recorded from M1 and Brodmann’s areas 3a and 2 exceeded this criterion (Figure 3B). In some cases, namely when comparing between single-predictor models in different coordinate frames, only the subset of neurons with multiple-predictor pseudo-*R^2^* > 0.20 were considered. This stricter threshold was chosen to avoid comparing single-predictor models in two coordinate frames that both performed poorly.

The number of joints in a neuron’s response field was determined as follows. First, because we incorporated different axes of rotation as separate predictors, and because we used both the posture and movement of each joint angle as a separate predictor, we grouped standardized regression weights according to the joints with which they were associated and calculated the sum of their squares. Once the sum of squared regression weights was computed for each joint, we determined the minimal set of joints that cumulatively explained 90% of their total sum.

The extent to which joints in a neuron’s response field (RF) were distributed across the entire hand was assessed by computing a co-occurrence matrix for each cortical field. For each pair of joints, we computed the number of neurons with *both* joints in their RFs, and normalized it by the number of neurons with either joint in their RFs.

We then used canonical correlation analysis to find the maximum correlation between each pair of joints — recall that individual joints can comprise separate predictors for different rotational degrees of freedom. We also determined the “proximity” of each pair of joints on the basis of the number of skeletal or ligamentous segments interposed between them. As a joint is defined as the junction between two or more bones (Table S2), two joints were deemed “adjacent” if both joints shared a bone or comprised adjacent metacarpal bones, the latter of which are connected by the transverse metacarpal ligament. This set of pairwise adjacencies formed an unweighted, undirected graph where nodes corresponded to joints and edges corresponded to links between adjacent joints. Minimum path lengths between all pairs of joints in this graph were then determined. Linear regression was then used to determine the extent to which the correlation or proximity of each pair of joints predicted the rates with which those joints co-occurred within neural RFs: the *R^2^* of this regression is reported.

### Interpreting encoding models: alternate coordinate frames

We assessed the degree to which two alternative kinematic coordinate frames might offer a more parsimonious description of neural activity than did joint angles: musculotendon lengths and principal components of joint angular kinematics. Musculotendon lengths were obtained from the same OpenSim model as the joint angles and yielded 35 different coordinates spread across 22 different muscles; multiple insertions of multi-articulate muscles were modeled as separate musculotendon units. Musculotendon lengths were only reconstructed for Monkeys 2-4. Joint principal component scores were obtained by applying PCA to joint angular kinematics on a monkey-by-monkey basis, pooling across sessions within each monkey but not across different animals. PCA was applied to joint angle kinematics, as has been done for previous attempts to reduce the dimensionality of hand kinematics (Thakur et al., 2008). In both cases, both the positions and velocities (derivatives) along the resultant kinematic dimensions were used to fit GLMs using methods similar to those used for models in a joint coordinate frame.

We also fit GLMs with a single predictor in each coordinate frame, seeking for each neuron the single joint DOF, single musculotendon DOF, and single principal component (PC), that maximized the pseudo-*R^2^* of the model fit. These models were fit at multiple latencies between neural and kinematic data, similar to the multi-predictor models. Pseudo-*R^2^* values were cross-validated using similar methods as for the multi-predictor models.

### Interpreting encoding models: testing for preferential encoding of posture or movement

Partial pseudo-*R^2^* of model *X* given model *Y* is computed using a calculation similar to that for partial coefficients of determination,

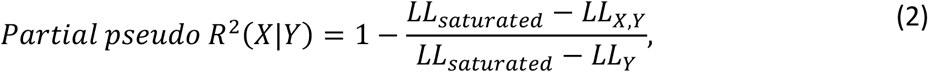

where LL_X,Y_ is the log likelihood of the combined model, and LL_Y_ is the log likelihood of the model using only the set of predictors *Y*. In essence, these computations quantify the amount of unique deviance explained by including predictors *X* after first removing all deviance that can be explained solely by predictors *Y*. As such, these partial pseudo-*R^2^* values are reported as “fraction of unique deviance explained” (FUDE), followed by the predictor set (either “posture” or “movement”) that filled the role of *X*.

### Testing spike history dependence and encoding of temporally extended hand trajectories

Previous reports of M1 neural encoding models of hand movements included spike history terms as predictors and used temporally-extended kinematic trajectories rather than instantaneous positions or velocities of joints (Saleh et al., 2010). Briefly, spike history terms were obtained by convolving spike trains with progressively wider causal filters, whereas temporally-extended kinematic trajectories were fit by treating each combination of kinematic degree of freedom and temporal lag as a separate predictor. In addition, a different parameterization of hand kinematics is used that incorporates fewer degrees of freedom, and a different measure is used to ascertain whether postures or movements are preferentially encoded. To determine the extent to which our results are dependent on such methodological differences (Figure S5), we fit models that implemented these approaches to determine at what point the interpretations of these models might differ.

