## Supplementary Material for "Postural Representations of the Hand in Primate Sensorimotor Cortex"

**Table S1.** Related to Figure 3. Statistics of pseudo-$R^2$ values of kinematic generalized linear models (GLMs). Statistics are shown for neurons with pseudo-$R^2 > 0.05$. For each dataset, we report the number of neurons above this criterion (N), the mean pseudo-$R^2$ across that population (μ), and its standard deviation (σ). The mean pseudo-$R^2$ in each dataset is compared to that of the reaching dataset using Welch's two-tailed *t*-test. The *t*-statistic (*t*), degrees of freedom (dof), and *p* value (*p*) are reported for each of these comparisons. Significant *p* values, determined via the Holm-Bonferroni method (FWER = 0.05) to correct for multiple comparisons, are highlighted in bold. *These particular GLM statistics are not reported in the original paper; rather, we fit GLMs using the approach described in the methods section but using neural and kinematic data from the Hatsopoulos et al. study. GLMs are similar to those used to predict firing rates of neurons during grasp, but instead of using 30-dimensional grasping kinematics to predict firing rates, we use 2-dimensional reaching kinematics (planar endpoint coordinates) to do so. All reported goodness-of-fit values are cross-validated as detailed in the *Methods*.

| *Task and Area* | *N* | *μ* | *σ* | *t* | *dof* | *p* |
|---|---|---|---|---|---|---|
| *Grasp Area 3a* | 32 | 0.201 | 0.120 | 2.92 | 54.4 | **5.02e-03** |
| *Grasp Area 2* | 36 | 0.149 | 0.096 | 1.07 | 73.0 | 2.89e-01 |
| *Grasp M1* | 167 | 0.211 | 0.148 | 4.74 | 118.4 | **6.11e-06** |
| *Reach M1 (Hatsopoulos et al 2007)** | 46 | 0.127 | 0.043 | | | |

**Table S2.** Related to Figure 5. List of non-stationary joints and muscles present in the musculoskeletal model of the arm. Inverse kinematics reconstructed the angle of each degree of freedom of each joint and the length of each muscle head of each muscle. *Not reconstructed for Monkey 1. **The symbol "X" stands in for one of digits 2-5 for this row of the table. ***Comprised 2, not 3, degrees of freedom in Monkey 1. †Use of human model leaves one fewer head than seen in the macaque, where these muscles are replaced by extensor digiti 2-3 and extensor digiti 4-5, respectively. ‡The fifth head that inserts onto the thumb in monkeys is absent in the human model; the flexor pollicis longus in the human model (a muscle that is absent in the monkey) is assumed to comprise this fifth head.

| *Joint* | *# DoF* | *Bones* | |
|---|---|---|---|
| *Elbow** | 1 | • Humerus<br>• Ulna | • Radius |
| *Wrist* | 3 | • Radius<br>• Carpus | • Ulna |
| *Carpo-metacarpal (CMC) 1* | 3*** | • Carpus | • Metacarpal (MC) 1 |
| *Metacarpo-phalangeal (MCP) 1* | 2 | • MC 1 | • Proximal phalanx (PP) 1 |
| *Interphalangeal (IP) 1** | 1 | • PP 1 | • Distal phalanx (DP) 1 |
| *CMC 4* | 1 | • Carpus | • MC 4 |
| *CMC 5* | 3*** | • Carpus | • MC 5 |
| *MCP X*** | 2 | • MC X | • PP X |
| *Proximal IP X*** | 1 | • PP X | • Middle phalanx (MP) X |
| *Distal IP X*,*** | 1 | • MP X | • Distal phalanx X |

| *Muscle** | *# Heads* | *Muscle** | *# Heads* |
|---|---|---|---|
| *Triceps brachii* | 3 | *Flexor carpi radialis* | 1 |
| *Biceps brachii* | 2 | *Flexor carpi ulnaris* | 1 |
| *Anconeus* | 1 | *Palmaris longus* | 1 |
| *Brachialis* | 1 | *Extensor digitorum communis* | 4 |
| *Brachioradialis* | 1 | *Extensor indicis proprius* | 1† |
| *Supinator* | 1 | *Extensor digiti minimi* | 1† |
| *Pronator teres* | 1 | *Flexor digitorum superficialis* | 4 |
| *Pronator quadratus* | 1 | *Flexor digitorum profundus* | 5‡ |
| *Extensor carpi radialis (ECR) longus* | 1 | *Extensor pollicis (EP) longus* | 1 |
| *ECR brevis* | 1 | *EP brevis* | 1 |
| *Extensor carpi ulnaris* | 1 | *Abductor pollicis longus* | 1 |

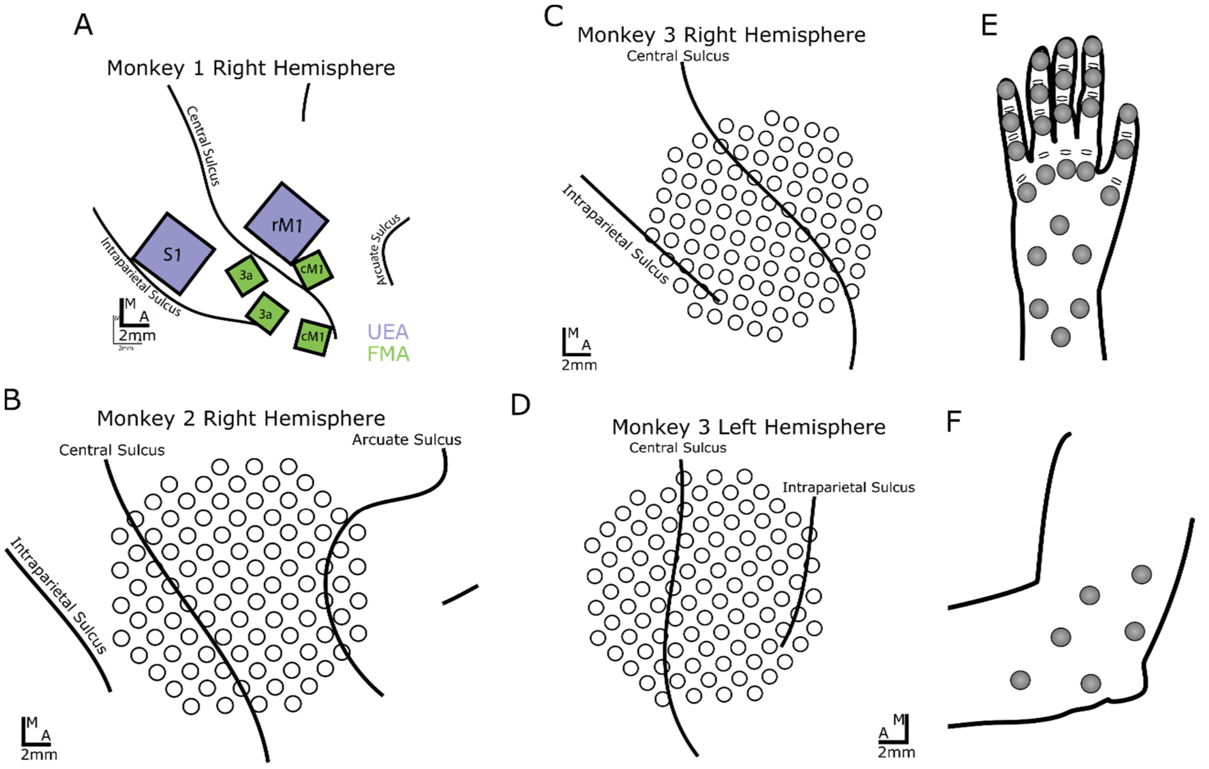


**Figure S1.** Related to Figure 1. Supplement to the Methods. Array placements relative to an oblique view of the cortical surface are shown for Monkey 1 **(A)**, Monkey 2 **(B)**, and the right **(C)** and left **(D)** hemispheres of Monkey 3. Figure 1 in the main text displays array placement in Monkey 4, in addition to histological reconstruction of architectonic borders and electrode locations. The placements of reflective markers relative to the hand **(E)** and elbow **(F)** are also shown.

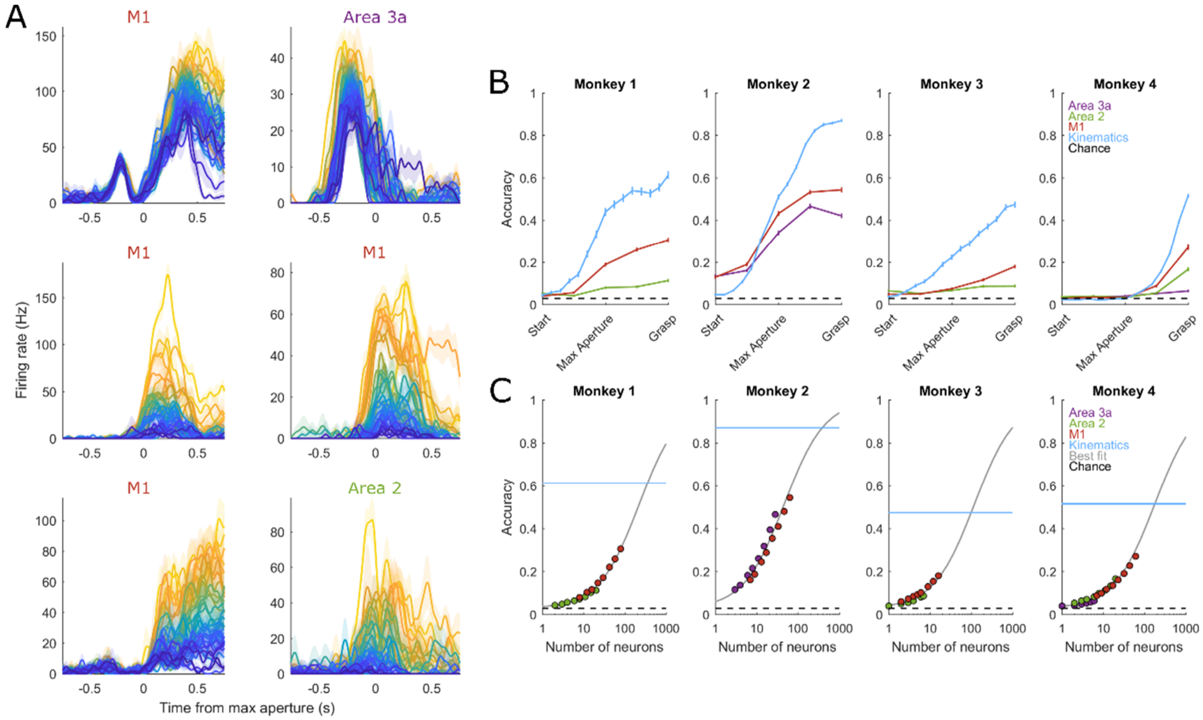


**Figure S2.** Related to Figure 2. Different objects give rise to a variety of different neural responses. **(A)** Example peri-event time histograms (PETHs) from 6 neurons across sensorimotor cortices that illustrate the variety of response profiles seen. PETHs are centered on maximum aperture. Different colors indicate different objects, ranked from weakest to strongest response on a neuron-by-neuron basis; colors from one PETH do not signify the same objects that they do in other PETHs. **(B)** Classification accuracy on the basis of neural population activity and hand kinematics for each of 4 monkeys. Vertical lines indicate ±1 S.E.M. Pseudo-populations were constructed on a monkey-by-monkey basis by aligning spike trains to each epoch and combining across recording sessions. Population sizes are different across cortical fields and monkeys. For Monkey 1: The area 2 pseudo-population comprises 20 neurons and the M1 pseudo-population comprises 80. For Monkey 2: Area 3a comprises 28 neurons and M1 comprises 63. For Monkey 3: Area 2 comprises 7 neurons and M1 comprises 16. For Monkey 4: Area 3a comprises 6 neurons, Area 2 comprises 20, and M1 comprises 61. Neural classification is based on spike counts integrated over a causal 500ms rectangular window. Neural classification accuracy surpasses chance performance in all cases in a manner that depends on population size and the amount of object information in each particular monkey's kinematics. Neural classification accuracy also follows a time course similar to that of kinematic classification performance, varying smoothly over the course of a trial. Neural classification accuracy exceeds chance prior to the start of movement in Monkey 2, which could be attributed in part to unmeasured isometric forces in anticipation of particular objects. **(C)** Peak classification accuracy as a function of population size. Logistic regression models ("Best fit") are fit separately for each monkey to account for their grasping strategies and, in turn, their different peak kinematic classification accuracies. In each monkey, extrapolation suggests that a few hundred neurons in each cortical field are sufficient to support object classification at the same level as the joint kinematics themselves.

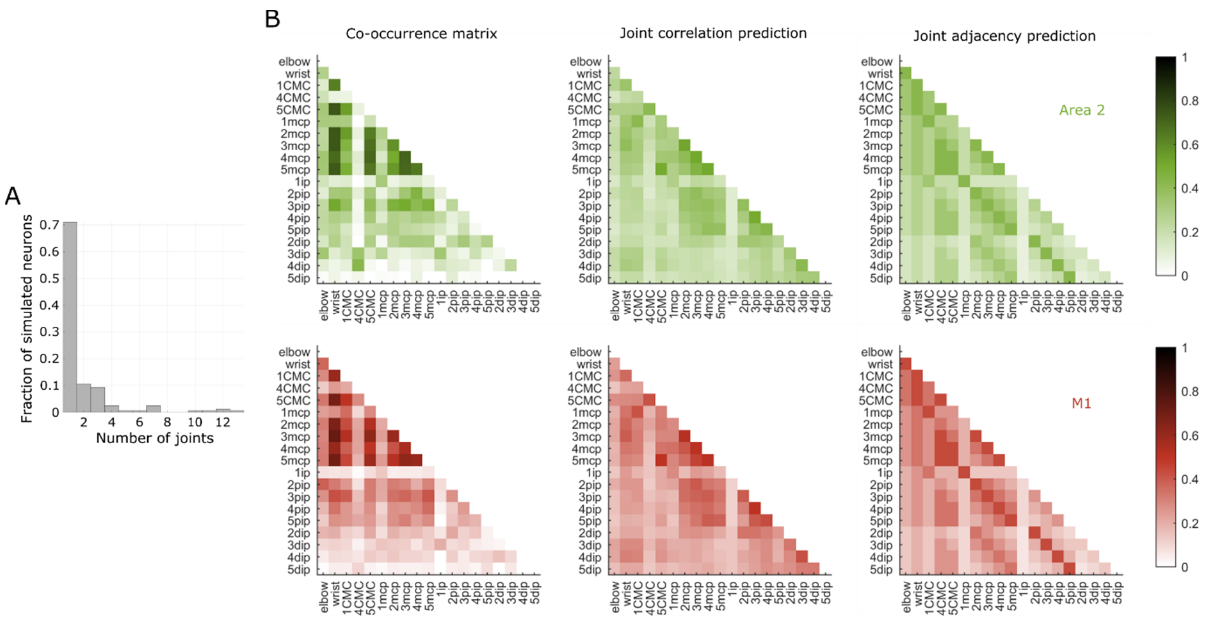


**Figure S3.** Related to Figures 4 and 5. Large RFs spanning the entire hand defy trivial explanation and are present in all sensorimotor cortical fields. **(A)** For simulated rate-varying Poisson neurons that track only one joint, a histogram of the number of joints in each simulation's response field (RF) inferred from GLM regression weights. Goodness-of-fit distributions are, by design, similar to what is observed in the neural data. Even with similar levels of noise relative to the neural data, the vast majority of simulations have GLM-inferred RFs containing just one joint. Thus, multi-joint RFs are unlikely to emerge from neurons that truly only track a single joint. **(B)** Similar to Figure 6 in the main text, but instead showing co-occurrence matrices across neurons in area 2 (*top row*) and M1 (*bottom row*). Just as in area 3a, a substantial fraction of variance is left unexplained in the joint correlation (Area 2: $R^2$ = 0.254; M1: $R^2$ = 0.277) and joint proximity (Area 2: $R^2$ = 0.146; M1: $R^2$ = 0.278) predictions of co-occurrence.

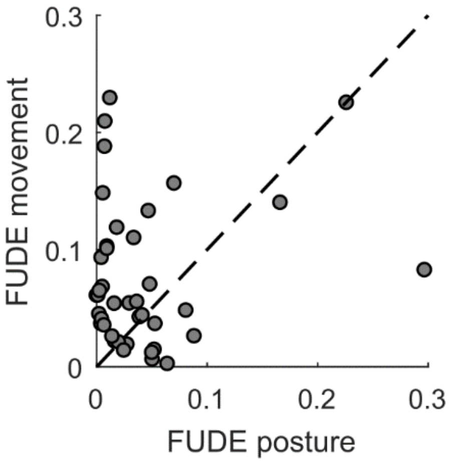


**Figure S4.** Related to Figure 7. Proximal limb M1 firing rates preferentially track limb movements rather than postures. Similar to Figure 7B, only showing the fraction of unique deviance explained (FUDE) by posture and movement models for M1 neurons during reaching (Hatsopoulos, Xu, & Amit 2007). The majority of neurons fall above the diagonal. In other words, more unique deviance is explained by limb movements than postures during reaching. This is consistent with previous reports (Paninski, Fellows, Hatsopoulos, & Donoghue 2004, Wang, Chan, Heldman, & Moran 2007), showing that the GLMs we use do not inherently perform better with postural predictors.

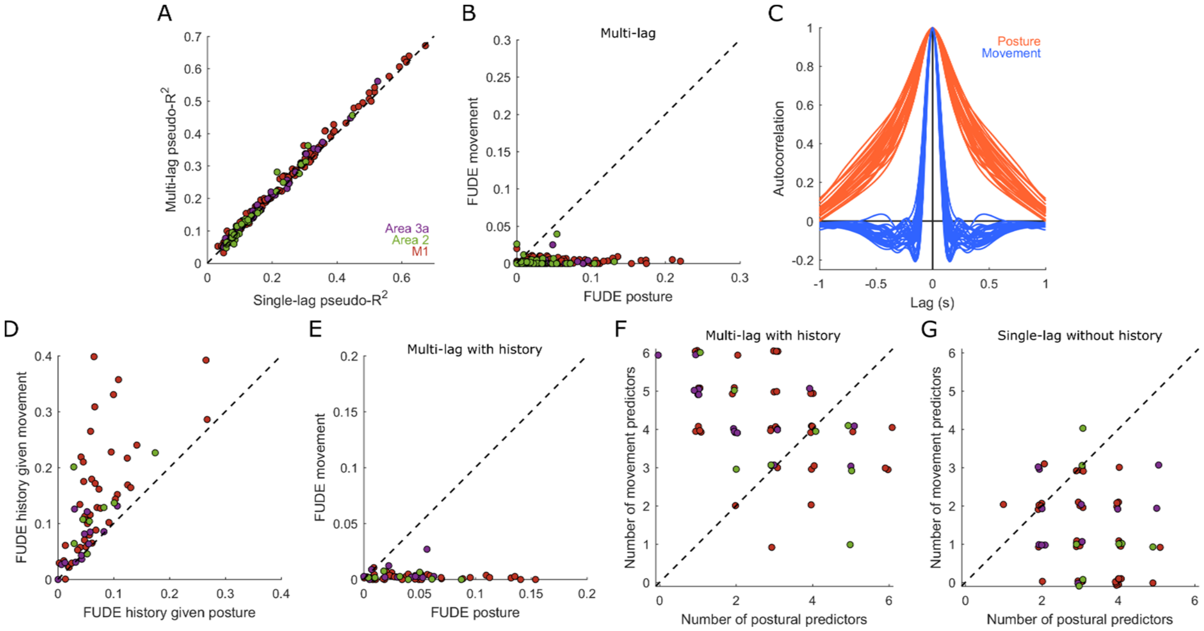


**Figure S5.** Related to Figure 7. Reconciling observed posture preference with past results that seem to show the opposite (Saleh et al., 2010). **(A)** Previous reports use GLMs that fit kinematic trajectories comprising multiple lags to the spiking activity of neurons. In the present study, we only use kinematics at a single optimal lag for ease of interpretation: Indeed, a multi-lag postural model supports the creation. We find that the improvement in goodness-of-fit (pseudo-$R^2$) when using predictors at multiple lags is minute: On average, a multi-lag model explains a single percentage point more in terms of the fraction of deviance explained by a single-lag model. Nonetheless, this difference is significant (paired-sample T-test, $t(234) = 13.17$, $p = 5.11\text{e-}30$, 95% confidence interval = [0.0089, 0.0121]). **(B)** However, even among multi-lag models, we find that the fraction of unique deviance (FUDE) explained by postural predictors (abscissa) greatly exceeds that of movement predictors (ordinate). **(C)** Mean autocorrelation functions of each of the 30 posture and movement degrees of freedom. Previous reports use GLMs that include spike history terms, whereas the present study does not. *S*pike history could preferentially affect deviance explained by postural predictors relative to that explained by movement predictors by virtue of *the* wider autocorrelations of the former. **(D)** The FUDE by spike history after accounting for hand posture predictors (abscissa) is indeed lower than that after accounting for hand movement predictors (ordinate), suggesting that the overlap between spike history *and posture* is greater than that between history and movement. **(E)** *E*ven when accounting for spike history, the FUDE by postural models (abscissa) still far exceeds that of movement models (ordinate). **(F)** Previous reports removed predictors one-by-one and counted the number of posture and movement predictors that resulted in a significant performance decrease. Using the 90%-of-squared-norm criterion (Figure 4B*)* to approximate this process*,* we find that most neurons' response fields (RFs) comprise more movement (ordinate) than posture (abscissa) degrees of freedom*,* at least when analyzing multi-lag models that include spike history. However, this contradicts the conclusion arising from the analysis of FUDE. **(G)** When analyzing single-lag models without spike history, this result is reversed. Neurons' RFs tend to comprise fewer movement (ordinate) than posture (abscissa) predictors, in line with what would be expected given the relative FUDE by the two. Note, the number of predictors is smaller here than in the main text because GLMs are fit to the reduced predictor set used in Saleh et al., 2010. We conclude that an interpretation that favors preferential encoding of postures over movements is robust to numerous methodological differences in constructing and analyzing neural encoding models, with only a particular method that confounds interpretation appearing to reveal preferential movement encoding.
